# Autophagy is suppressed in peripheral blood mononuclear cells during chronic obstructive pulmonary disease

**DOI:** 10.1101/2024.04.27.591479

**Authors:** James M Cooper, Shiye Chen, Susan E Lester, Julia Kim, Jason Gummow, Thomas Crowhurst, Emily Lawton, Arash Badiei, Phan T Nguyen, Paul N Reynolds, Hubertus PA Jersmann, Eugene Roscioli

## Abstract

Assessing autophagy may offer insights into the pathogenesis of chronic obstructive pulmonary disease (COPD). However, measuring the dynamic aspect of autophagy is challenging, and sample manipulation can cause signal fluctuations that deviate from physiological conditions. We applied an organotypic method to quantify autophagy in COPD, where it frequently demonstrates disease-related dysregulation. Blood from control and COPD participants were treated with or without chloroquine. LC3B-II abundance was quantified in peripheral blood mononuclear cells, and findings were validated by transmission electron microscopy. Our observations show that while basal LC3B-II abundance was similar between groups (P = 0.60), autophagic flux was significantly lower in the COPD cohort, suggesting disruption in the regulatory factors that direct autophagosome clearance (P = 0.004). This was supported by less frequent observations of autophagy-related vacuoles in the cytosol of COPD-derived PBMCs. Our findings indicate that the suppression of autophagy can be detected in the blood of individuals with COPD, which warrants further investigation into its contribution to extrapulmonary disease processes.

## Introduction

Chronic obstructive pulmonary disease (COPD) is among the group of conditions in which dysregulated macroautophagy (hereafter referred to as autophagy) is understood to potentiate disease and may contribute to causation. Many disease phenotypes seen in COPD, including the autoimmune-like inflammation [1], persistent infection [2], and accelerated senescence [3], align with the consequences of restricted autophagic flux. Despite these findings, there remains an important debate to determine whether autophagy is restricted, and thereby diminishing its homeostatic support, or accelerated as a maladaptive response to persistent COPD-related exposures (reviewed in [4]). In broad terms, this uncertainty is because COPD is complex, has a genetic component, and is typically diagnosed at an advanced stage when the convergence of several pathophysiological events obscure causal molecular patterns. Coupled with this, assessing autophagy function in a manner that reliably reflects the situation *in vivo* remains challenging [5]. Hence, despite the significant promise of targeting autophagy to manage COPD, the world’s third most lethal disease [6], autophagy remains largely disconnected from clinical, diagnostic and pharmacological inquiry.

Addressing the measurement problem, a recent investigation has described an organotypic method to assess autophagy function in peripheral blood mononuclear cells (**PBMC**), by directly treating blood samples with/out chloroquine (**CQ**) and assessing readouts of LC3B-II [7]. The value of this method is that the *ex vivo* measures of autophagic flux approximate the situation in the participant’s PBMCs; taking blood requires minimal clinical resource, the period from collection to protein isolation is brief, and the blood maintains the PBMCs as if they were still in the body. While the lethal processes ascribed to COPD occur in the airways, this condition is associated with serious systemic effects [8]. The products of inflammation and inhaled environmental factors (e.g. chemicals from tobacco and occupational/environmental exposures), are transmitted to cells of the circulatory system [9, 10]. Further to this, disease-associated genetic variants are present in all somatic cells, meaning that immune cells encoding COPD susceptibility alleles may be affected both by pulmonary-derived exposures and independently through intrinsic mechanisms. Indeed, the pleiotropic nature of many COPD risk variants, capable of influencing both pulmonary and systemic traits such as metabolic syndrome and inflammation, is supported by genome-wide studies that often source genetic material from non-airway tissues [11]. Autophagy-related genes are linked (or are convincing candidates) to the development of human disease, including COPD, asthma and pulmonary fibrosis [12], and we are now identifying the relationship between the peripheral consequences of COPD and dysregulation of autophagy [13-15]. Hence, an interesting question that arises from these findings is whether assessing autophagy in the peripheral circulation can inform the development of a non-invasive screen for COPD.

Here we assessed autophagy in PBMCs isolated from individuals suffering from COPD, to evaluate alterations in autophagic flux. Our observations show that PBMCs from participants with COPD exhibits arrested autophagy compared to the outcomes from healthy controls.

## Results

### PBMCs from COPD participants exhibited reduced autophagic flux

The observations for LC3B-II protein abundance in PBMCs were first compared separately for each group/exposure (**Figure 1A-B**; participant demographics are shown in **Table 1**). PBMCs from either group exhibited a similar near-zero signal distribution for the untreated condition (P = 0.60, control vs COPD), indicating low basal LC3B-I lipidation for PBMCs is normal for these groups. Both control and COPD cells also demonstrated an elevation in LC3B-II abundance after exposure to CQ (both P < 0.001, CQ vs NT). However, control PBMCs accumulated an appreciably larger amount of LC3B-II in response to CQ over the exposure interval (P < 0.007, control vs COPD), suggesting that PBMCs from COPD participants exhibit restriction in the processes contributing to autophagic flux (see also **Figure S1**). Values for autophagic flux (△ LC3B-II) were derived from the difference between LC3B-II abundance in CQ-treated and untreated samples, reflecting LC3B-II accumulation following lysosomal inhibition. Shown in **Figure 1C**, the COPD group exhibited significant autophagic insufficiency when compared to the control group (P = 0.004, control vs COPD). The observations for autophagic flux were further assessed to determine whether differences in age and gender were significant confounding variables (model-based analysis). Shown in **Table 2**, the reduction in autophagic flux determined for the COPD group remained significant after accounting for the covariates. However, estimates derived within the model show that age (gender was a non-significant confounder), can influence outcomes related to autophagic flux. The efficacy/specificity of the LC3B-I/II antibody was also assessed using lentivirus CRISPR KO directed to *ATG5*. Depletion of ATG5 protein caused a reduction in ATG5-mediated lipidation of LC3B-I (i.e. a reduction in LC3B-II), concomitant with an elevated abundance of Sequestosome-1 (**Figure 1D**).

**Figure 1.**
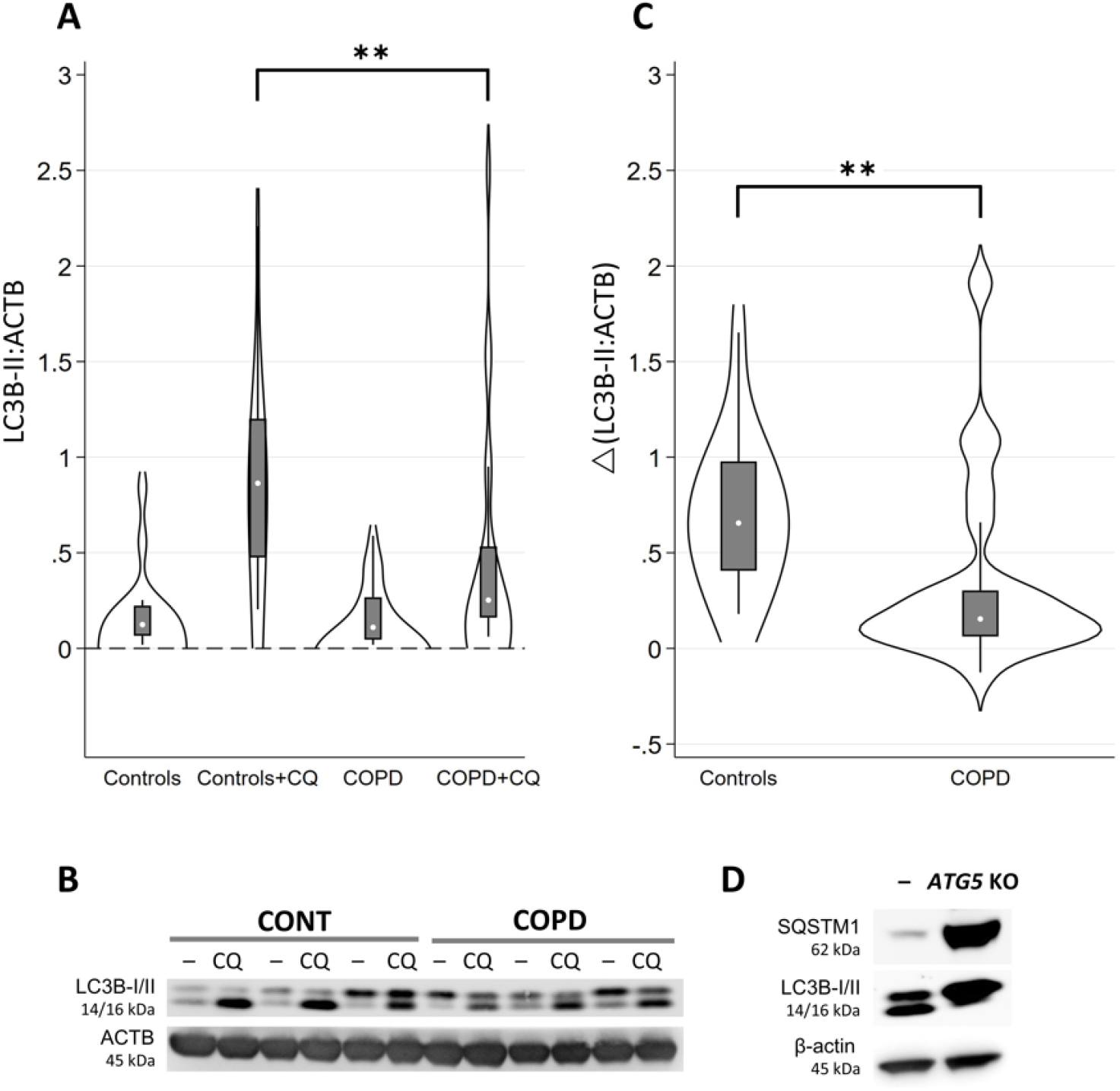
Peripheral blood mononuclear cells from individuals with COPD exhibited reduced autophagic flux. (**A**) Basal level LC3B-II observation densities show a near-zero distribution for both groups, while the COPD group exhibited a significant reduction in LC3B-II accumulation with chloroquine (CQ), vs the controls. (**B**) A representative blot comparing no-treatment and CQ outcomes for LC3B-I/II and Actin-β (ACTB, **C**). Measures of autophagic flux (△ LC3B-II) showed a significant reduction in autophagy competency for the COPD group. (**D**) The LC3B-I/II antibody was validated using protein isolated from 16HBE14o-cells transduced with a CRISPR ATG5 KO construct. The depletion of ATG5 protein reduced detectable LC3B-II (the lipidated form of LC3B-I), and substantial increases in the LC3B-I and p62/Sequestosome-1 (SQSTM1) signals. Outcomes for LC3B-II were normalized to the Actin-β signal. Control, n=15 and COPD, n=22. “**” denoted P < 0.01. Outcomes for the violin graphs were from non-parametric analyses; Wilcoxon matched-pairs sign-rank test when comparing no-treatment and CQ, and Kruskal-Wallis rank test for comparisons between control vs COPD. Violin graphs boxplots show the mean ± standard error and whiskers denote ± 95% CI.

**Table 1:**
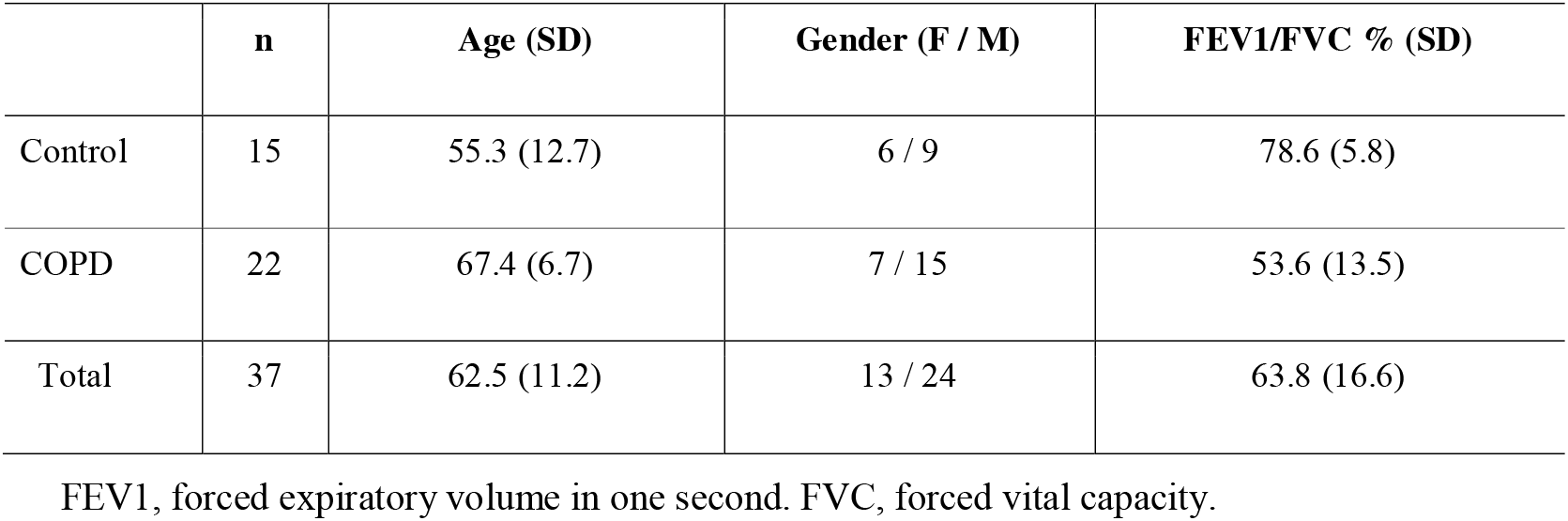
Participant demographics for protein biochemistry.

**Table 2:** Model-based analysis of autophagic flux adjusted for age and gender.

| Covariates | ΔLC3B-II:Actin-β (95% CI) | | COPD vs Controls |
| --- | --- | --- | --- |
|  | COPD | Controls | P value |
| None | 0.37 (-0.01, 0.76) | 0.82 (0.42, 1.23) | <0.0001 |
| Age, Sex <sup>1</sup> | 0.35 (-0.03, 0.73) | 0.59 (0.24, 0.94) | 0.003 |

<sup>1</sup>Estimated at the covariate distribution of COPD patients i.e. the predicted values if controls had been age/sex matched to COPD patients.

### Autophagosome abundance is decreased in PBMCs exposed to CQ in COPD vs control

High sensitivity microscopy was performed to directly inspect PBMCs for alterations in autophagosome frequency between the control and COPD conditions. Shown in **Figure 2 A-B**, vesicular structures indicative of autophagy-related vacuoles were rarely detected in either control or COPD-derived PBMCs. However, the accumulation of autophagy-related vacuoles were increased in control PBMCs challenged with CQ. In conjunction with the outcomes from protein biochemistry, these observations are consistent with reduced autophagic flux in PBMCs obtained from individuals with COPD.

**Figure 2.**
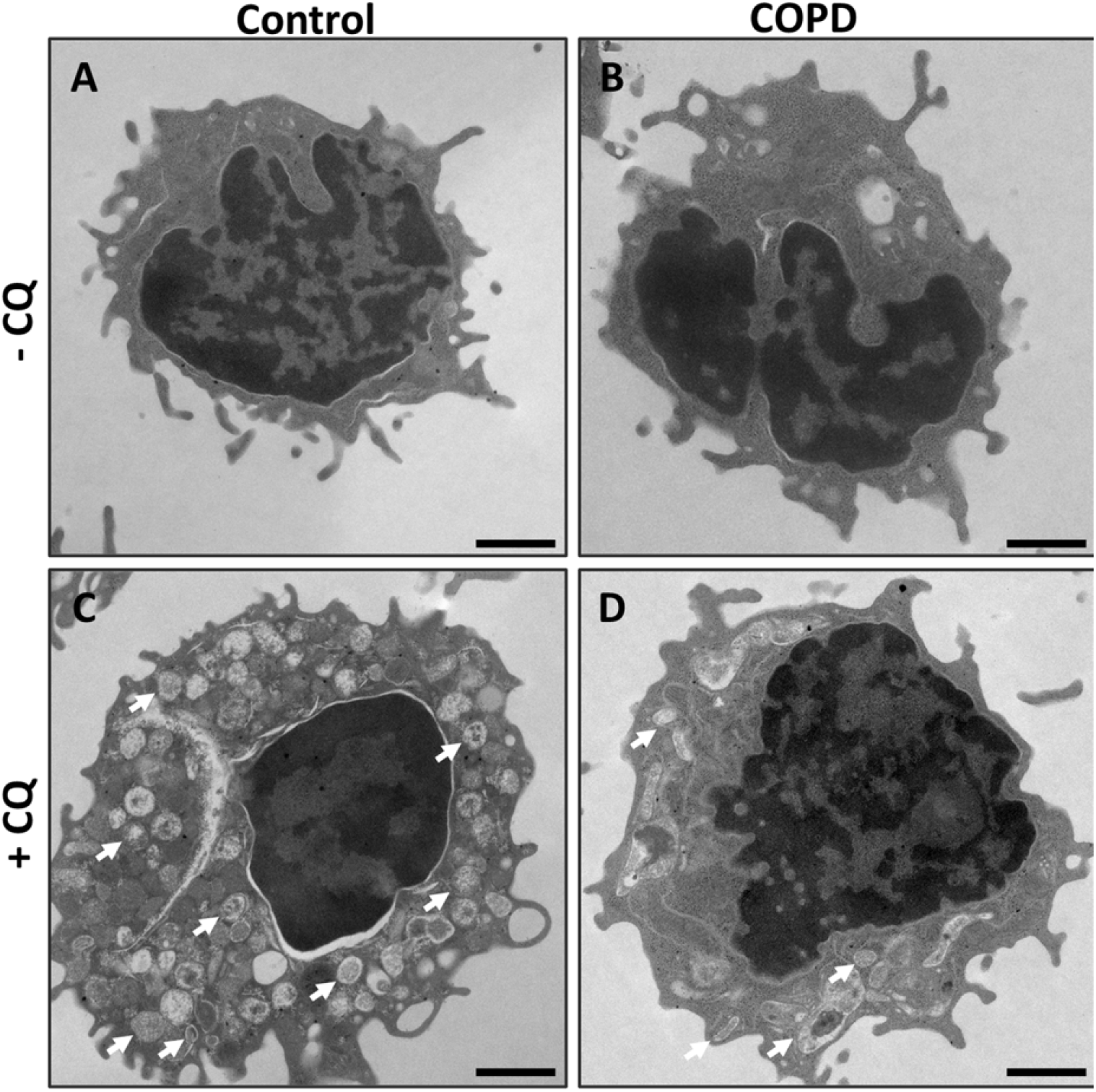
Autophagy-related vacuoles accumulate in PBMCs following chloroquine exposure. (**A-D**) Representative transmission electron micrographs of untreated peripheral blood mononuclear cells (PBMCs) isolated from control and COPD participants or following exposure to chloroquine (CQ; 150 μM, 1 h). (**A-B**) PBMCs from control and COPD participants under untreated conditions showed low basal levels of autophagy, with autophagy-related vacuoles rarely detected. (**C-D**) Exposure to CQ resulted in the accumulation of autophagy-related vacuoles (examples indicated by white arrows). Consistent with the biochemical flux measurements shown in Figure 1, vacuoles appear more abundant in control PBMCs compared with COPD-derived PBMCs. Scale bars: 1 µm.

## Methods

### Participants

Ethics approval was granted by the Central Adelaide Local Health Network Human Research Ethics Committee (CALHN reference number: 12978). All participants provided written informed consent. Samples were obtained by the clinical team at the Royal Adelaide Hospital’s Department of Thoracic Medicine and Queen Elizabeth Hospital’s Respiratory Medicine Unit. Participant demographics are shown in **Table 1**. Control participants reported no history of chronic respiratory disease and were never smokers. COPD was determined by clinical observations and lung function test (FEV1/FVC ratio below 70%).

### PBMC isolation

PBMCs were isolated without deviation in processes and products described by Bensalem et al. (2020) [7]. Briefly, blood collected in lithium-heparin tubes were entered into the PBMC isolation process no longer than 2 hours after obtaining the sample from the clinic. Chloroquine diphosphate was added to a concentration of 150 _µ_M and allowed to incubate for 1 h at 37°C, simultaneously with the untreated sample. PBMCs were isolated using the Ficoll gradient method.

### Western blot

Western blots were performed as previously described (e.g. [16]). Briefly, protein was isolated using M-PER Mammalian Protein Extraction Reagent (Thermo Fisher Scientific Inc, 78501), and Halt Protease and Phosphatase Inhibitor Cocktail (Thermo Fisher Scientific Inc, 78441). Protein was quantified using the Pierce BCA Protein Assay Kit (Thermo Fisher Scientific Inc, 23225), and 10 μg electrophoresed in 4–12% Bis-Tris denaturing gels (Thermo Fisher Scientific, NP0321) using MOPS chemistry (Thermo Fisher Scientific, NP0001). When possible, a calibrator protein sample of known concentration was included to account for signal variation due to inter-experimental/membrane variation. Transfer was to 0.2 µm pore PVDF membranes (Bio-Rad Laboratories Inc, 1704156), and sectioned so target and normalization signals could be quantified from the same blot to mitigate loading inconsistencies, inter-membrane variation and detection bias caused by multi-probing or strip/re-probing the membrane. Membranes were blocked in 5% skim milk for mouse anti-Actin-β (1:10,000, A1978; Sigma-Aldrich Co.), and 5% BSA (Merck, 126575), for rabbit anti-LC3B-I/II (Cell Signaling Technology, 4108) and subsequently probed overnight at 4°C with the same respective buffer/antibody combinations. Secondary incubation was 1 hour at RT in 5% skim milk with mouse or rabbit IgG horseradish peroxidase-conjugated antibodies (R&D Systems, HAF007 and HAF008, respectively). The suspension for blocking and antibody incubations was 1x Tris buffered saline (Thermo Fisher Scientific, J60877.K3), with 0.1% Tween 20 (Merck, P1379). Markers were sectioned off prior to imaging to prevent primary/secondary antibody cross-reaction/non-specific binding. Chemiluminescent signal production was with Amersham ECL Advanced Western Blotting Detection reagents (GE Healthcare, RPN2135). Image acquisition was performed using the LAS-4000 Luminescent Image Analyzer (Fujifilm Life Sciences, Japan), with signal detection set to annotate blots that produced signals beyond a predetermined linear range for each target/probe. Histogram densitometry was performed using Multi Gauge software (Fujifilm Life Sciences, V3.1, Japan).

### Generation of LentiCRISPRv2-ATG5

Briefly, HEK-293T cells were co-transfected with lentivirus construct encoding *Cas9* and a sgRNA targeting exon 7 of *ATG5* (LentiCRISPRv2-*ATG5*; Addgene, 99573 [17]; deposited by Edward Campbell), pMDLg/pRRE (Addgene, 12251), pMD2.G (Addgene, 12259), and pRSV-REV (Addgene, 12253) (Packaging vectors deposited by Didier Trono) using Lipofectamine 3000 (ThermoFisher Scientific, L3000150). Supernatant was collected 24 and 48 hours after transfection, centrifuged at 3000 rpm, filtered (Corning, 430768), and ultra-centrifuged (18000 rpm; ThermoFisher Scientific, Sorvall WX80). After ultra-centrifugation, supernatant was removed and the pellet resuspended and preserved at -80°C. Titration was performed by transducing HEK293T cells in a 24-well plate with LentiCRISPRv2-*ATG5*, and after 72 hours transduction the cells were collected and DNA extracted/purified. DNA for cells transduced with Lenti GFP was used as a standard ladder and qPCR was performed and normalized to house-keeper gene to determine LentiCRISPRv2-*ATG5* titre.

### Generation of ATG5 KO 16HBE14o-cells

The LC3B-I/II antibody applied here is known to be sensitive and specific for LC3B-II [18]. To verify this, 16HBE14o-cells (EMD Millipore Corporation, SCC150), were used to model the functional knockout of autophagy. This cell line was chosen as 16HBE14o-produced high LC3B-1I and low Sequestosome-1 signals during log phase growth (i.e. to allow sensitive comparison vs outcomes for *ATG5* KO), and to avoid applying clinical samples (i.e. PBMCs), to optimization procedures. 16HBE14o-cells (EMD Millipore Corporation, SCC150), were propagated according to the manufacturer’s protocol and transduced with LentiCRISPRv2-*ATG5* at an MOI of 10 and allowed to culture for 48 hours. Selective pressure on transduced cells was applied by culturing in the presence puromycin (ThermoFisher Scientific, J67236.XF) for 15 days, after which colonies were selected, propagated and screened. To confirm ATG5 protein knockout, western blot analysis was performed using anti-rabbit-ATG5 (1:2000; AbCam, Ab108327). Functional inhibition of autophagy was assessed by Western blot using LC3B-I/II (as above) and Sequestosome-1 (Cell Signaling Technology, 5114) antibodies.

### Transmission electron microscopy

PBMC isolates were pelleted (400 g) for 5 minutes and resuspended/fixed in 500 µl of TEM fixative (4% formaldehyde, 1% glutaraldehyde, 4% sucrose in 1x PBS; each Sigma-Aldrich). Fixed cells were processed for mounting by the staff at the University of Adelaide Microscopy suite. Sections were examined using an FEI Tecnai G2 Spirit transmission electron microscope (Thermo Scientific, Eindhoven, Netherlands). Autophagic vacuoles were identified as vesicular structures containing heterogeneous electron-dense cytoplasmic or membranous material and bounded by a limiting membrane. Vesicles that were completely electron-lucent or displayed uniform electron density consistent with lysosomes were excluded from quantification.

### Statistical analysis

For each protein, Western blot densitometry results were analyzed with a random effect of test repetition and reported relative to the abundance of Actin-β. Analysis was performed in Stata v18 (StataCorp LLC). LC3B-II western blot intensities were analyzed using a gamma (log link) multi-level regression model, with log(Actin-β) intensities as an offset and a robust variance-covariance. Random effects included individuals nested within blot, and CQ treatment as a crossed random factor for each individual. Fixed effects included group (COPD vs control), CQ treatment (yes vs no) and the group × CQ interaction. Postestimation tools were used to estimate marginal means for LC3B-II normalized to Actin-β (LC3B-II:Actin-β), and the difference in these marginal means due to CQ treatment (LC3B-II:Actin-β). Age and sex were included as additional covariates, and a full factorial model was estimated. The marginal means for both COPD and controls were then estimated using the age/sex covariate distribution of COPD participants i.e. the results for the controls were estimated according to the age/sex distribution as COPD participants to allow a direct comparison.

## Discussion/Conclusion

We show that autophagic flux is reduced in PBMCs from individuals with COPD using an organotypic CQ flux assay. This method afforded a temporal window to assess autophagy in an organotypic manner (i.e. *ex vivo* exposure to CQ), which enabled us to quantify and compare the relative magnitude of autophagic function available to PBMCs from control and COPD participants. Comparative reports for COPD and related conditions that exhibit significant systemic inflammation, for the most part, explain the involvement of autophagy in disease processes with snapshots of LC3B-II and, although not always, an autophagy-receptor substrate (e.g. [19-22]). This can provide important information, but are not sufficient to describe how disease-related phenomena connect to the perturbations in autophagic competency. As flux implies a process (i.e. with a start and endpoint), it is important to consider the dynamic aspect of autophagy to provide information pertaining to function. Further to this, and important for PBMCs, measures of autophagy receptor abundance do not always correlate with the outcomes for autophagic flux inferred by the abundance of LC3B-II (e.g. [7, 23, 24]). Indeed, we tested three autophagy receptor proteins (NBR1, TAX1BP1 and p62/Sequestosome-1), in addition to phospho-ATG16L1 which is reported to correlate with autophagy induction [25], and neither provided outcomes that were consistent with the LC3B-II observations in PBMCs (**Figure S2**). This issue has been observed by several others (e.g. [7] for PBMCs), and underscores the challenges that need to be considered when assessing autophagy in a meaningful way. Undoubtedly, these measurement complexities are central to the current debate as to whether autophagy is inappropriately attenuated, or increased as a maladaptive response to COPD-related disease phenomena. Our findings, at least for PBMCs, point to a scenario whereby those COPD-related phenotypes that can be logically aligned with autophagic dysregulation, arise from inappropriate restriction of autophagic flux.

The utility and limitations of these outcomes can be addressed in terms of whether measures of autophagy can be used as a prospective diagnostic readout for COPD. While assessing blood is an attractive non-invasive tissue for diagnostic purposes, a central question is whether assessing autophagy in PBMCs is a suitable proxy for the airways. Despite our findings and the increasing evidence that peripheral lymphocytes are altered in COPD [10, 26], the vast airway epithelial and macrophage interface is the first to respond to most COPD-related environmental and microbial exposures. Hence, a rational position is that the assessment of lung tissues should offer more informative predictive insights, particularly during early disease. Indeed, outcomes derived from lung samples have been convincingly shown to exhibit autophagy incompetency in COPD [2, 27, 28]. Further, the abundance of autophagy receptors (to support LC3B-II observations) in e.g. airway epithelial cells, generally correlate with LC3B-II levels, meaning that their analysis can provide a robust assessment of autophagic flux [2]. Indeed, efforts in our studies using airway biopsies remain limited by blood and mucin co-contamination (e.g. airway brushing biopsies) which produces significant signal deterioration. To address this, we are currently developing/optimizing a model to reassess airway samples using an adapted version of the method applied here. In terms of achieving a balance of disease/control participant demographics, our findings and observations from other groups support the position that chronological age informs (and is influenced by) the state of the biochemical processes that regulate autophagy (**Table 1**; [27, 29, 30]). This is demonstrated in COPD, where accelerated senescence and unscheduled apoptosis are hallmarks of disease and are associated with the dysregulation of autophagy [3, 31-33]. While our evaluation of autophagy remained significant after correcting for the age disparity, assessments using homogeneous participant groups will be a central prerequisite leading to a diagnostic model that is predictive of COPD. Further, an overarching limitation of this approach is the reliance on Western blotting, a method that is labour-intensive, susceptible to interexperimental variation, and is a semiquantitative assessment of protein abundance. These limitations are particularly relevant in autophagy studies, where accurate interpretation requires additional validation to distinguish between increased flux and impaired autophagosome degradation.

We provide evidence that autophagy is suppressed in PBMCs from COPD donors. A clear understanding of the complex nexus between autophagy and COPD, particularly in a clinically relevant context, awaits the development of more efficient analytical methods yielding explicit outcome measures. Until then, a critical goal in this area of research will be to clarify whether autophagy dysregulation in the inflamed airway is a driver of COPD pathogenesis or merely a correlative phenomenon.

## Supplementary Materials

The following supporting information can be downloaded at: Figure S1: Additional graphical presentation of chloroquine-induced elevation of LC3B-II in peripheral blood mononuclear cells in control vs COPD.; Figure S2: Additional examination of autophagy-related markers.

## Author Contributions

Conceptualization, E.R.; methodology, E.R., J.M.C., S.C., S.E.L., J.K. and J.G.; investigation, E.R., J.M.C., S.C., S.E.L., J.K. and J.G.; resources, E.R., J.K., T.C., E.L., A.B., P.T.N., P.N.R. and H.P.A.J.; data curation, S.E.L., J.M.C., S.C., J.K. and E.R.; formal analysis, S.E.L.; writing-original draft preparation, E.R., J.M.C. and S.C.; writing-review and editing, all authors; project administration, E.R.; supervision, E.R.; funding acquisition, E.R. and H.P.A.J. All authors have read and agreed to the published version of the manuscript.

## Institutional Review Board Statement

This study was conducted in accordance with the Declaration of Helsinki and approved by the Human Research Ethics Committee of the Central Adelaide Local Health Network (CALHN) (approval number: 12978; approval date: 22 April 2021).

## Informed Consent Statement

Informed consent was obtained from all subjects involved in the study.

## Data Availability Statement

The data presented in this study are available upon request from the corresponding author.

## Acknowledgements

We are grateful for the expertise and support of the clinical staff of the Royal Adelaide Hospital Thoracic Department and The Queen Elizabeth Hospital Respiratory Unit, and the participants enrolled into this study who generously provided their valuable time and tissue samples. We are also grateful to the Adelaide Microscopy team for their expertise and support with electron microscopy.

---

**Figure S1.**
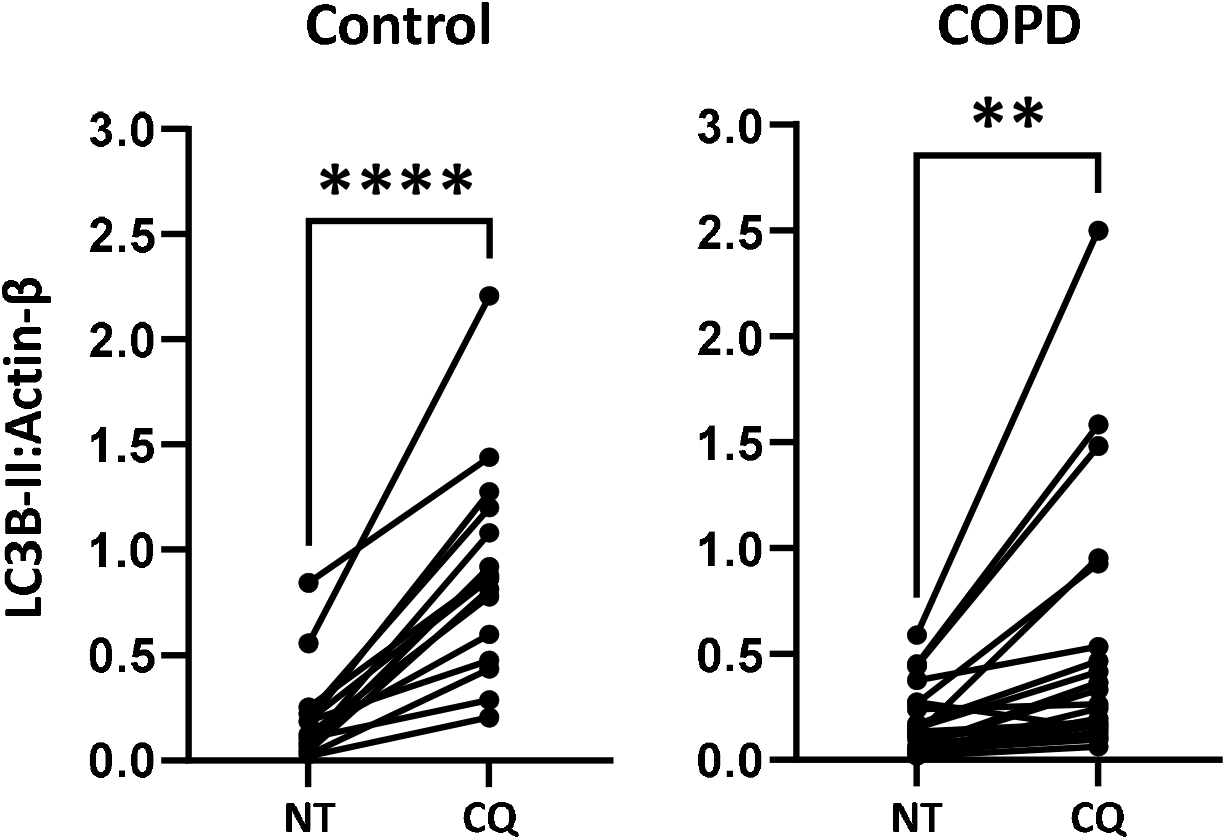
Chloroquine-induced elevation of LC3B-II in peripheral blood mononuclear cells is heightened in control vs COPD. Accumulation of the LC3B-II signal is more pronounced in the control group suggesting heightened autophagosome biosynthesis in the control group during the exposure period. Normality was confirmed (e.g. Shapiro-Wilk and D’Agostino–Pearson tests) and analysis was performed using paired t-tests. “**” denotes P < 0.01, and “****” P < 0.0001.

**Figure S2.**
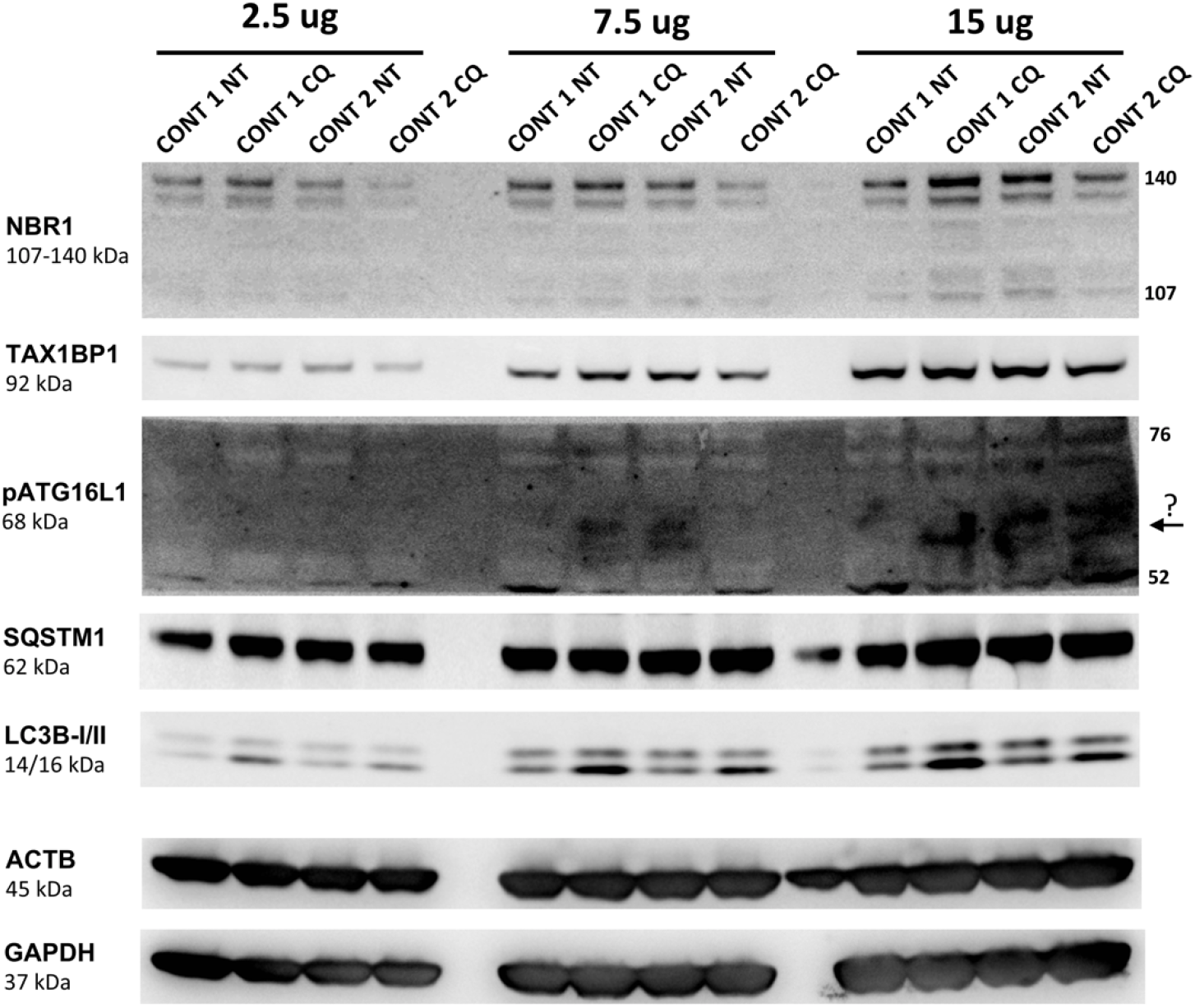
Autophagy receptors and pATG16L1 did not exhibit signal modulation in a manner consistent with the inhibition of autophagy in PBMCs exposed to chloroquine. Consistently, Western blot outcomes showed the autophagy receptors Neighbor of Brca1 (NBR1), TAX1BP1 and p62/Sequestosome-1 (SQSTM1), failed to demonstrate a concomitant signal induction with LC3B-II in PBMCs exposed to chloroquine (CQ; 150 µM for 1 hour vs no-treatment control; NT). Assessment of pATG16L1 is reported to quantify autophagy flux independent of LC3 or autophagy receptor proteins, was below the detection threshold in PBMCs (normalized transcripts per million values are approximately 2.1, vs e.g. Actin-β (ACTB) with approximately 1,000 nTPM (Proteinatlas.org)). This issue was compounded by the specified blocking agent being restricted for importation into Australia (personal communication with AbCam). Images are representative of several efforts to optimize these outcomes and are from a single membrane to mitigate inter-membrane variability and detection bias caused by multi-probing or strip/re-probing the membrane. Protein loading was 2.5, 7.5 and 15 µg to detect signal modulation that may be obscured by high signal magnitude (e.g. high protein abundance) or low signal strength (e.g. low abundance or low efficacy antibody-target interactions). Two normalizing proteins were used to account for signal modulation in either. NBR1 has been shown to migrate at either 140 or/and 107 kDa. Markers are removed to prevent primary/secondary antibody cross-reaction/non-specific binding.

